# Investigation of layer specific BOLD in the human visual cortex during visual attention

**DOI:** 10.1101/2021.02.07.430129

**Authors:** Tim van Mourik, Peter J. Koopmans, Lauren J. Bains, David G. Norris, Janneke F.M. Jehee

## Abstract

Directing spatial attention towards a particular stimulus location enhances cortical responses at corresponding regions in cortex. How attention modulates the laminar response profile within the attended region, however, remains unclear. In this paper, we use high field (7T) fMRI to investigate the effects of attention on laminar activity profiles in areas V1-V3; both when a stimulus was presented to the observer, and in the absence of visual stimulation. Replicating previous findings, we find robust increases in the overall BOLD response for attended regions in cortex, both with and without visual stimulation. When analyzing the BOLD response across the individual layers in visual cortex, we observed no evidence for laminar-specific differentiation with attention. We offer several potential explanations for these results, including theoretical, methodological and technical reasons. Additionally, we provide all data and pipelines openly, in order to promote analytic consistency across layer-specific studies, improve reproducibility, and decrease the false positive rate as a result of analytical flexibility.

## Introduction

Directing visual attention to a location in the visual field typically improves behavioral sensitivity to stimuli presented at that location (***Posner, 1980***; ***Lee et al., 1997a***; ***Yeshurun and Carrasco, 1998***; ***Carrasco et al., 2004***; ***Baldassi and Verghese, 2005***; ***Ling et al., 2009***). It is well known that these attentional benefits in behavior are accompanied by increases in BOLD response in early visual areas (e.g. ***Brefczynski and DeYoe*** (***1999***); ***Gandhi et al.*** (***1999***); ***Kastner et al.*** (***1999***)), but how top-down processes modulate cortical responses at the laminar level remains unknown.

It is known from anatomical studies that the human cerebral cortex can be subdivided into histological layers with different cell types. The cytoarchitectonic structure varies across the brain and forms the basis of the Brodmann atlas (***Brodmann, 1909***). While the precise function of each cortical layer remains unclear, their connectivity profile suggests a division in terms of bottom-up and top-down processing (***Felleman and Van Essen, 1991***; ***Barone et al., 2000***; ***Shipp, 2016***). Most brain areas have six different histological layers. Specifically, Layer IV and to a lesser extent Layer V/VI are commonly associated with receiving feedforward drive from Layer III of lower cortical areas or from the thalamus (***Jones, 1998***; ***Constantinople and Bruno, 2013***). Layers I-II and VI, in contrast, are typically implicated in receiving downward information flow (feedback), which often originates from layer V (***Alitto and Usrey, 2003***). This bottom-up versus top-down connectivity profile of each of the layers is to some degree also paralleled in functional data. That is, from neurophysiological and neuroimaging work, it is known that various visual stimuli and tasks can exert differential effects on the various layers (***Maier et al., 2010***; ***Xing et al., 2012***; ***Self et al., 2013***; ***Vélez-Fort et al., 2014***; ***O’Herron et al., 2016***). Intracranial work in monkeys, for instance, shows that for selective attention and working memory (two functions that are commonly associated with top-down processes), current source density is increased in deep and superficial compared to middle layers in primary visual cortex (***van Kerkoerle et al., 2017***). Similar layer specific patterns have been shown in animal functional MRI. For instance, whisker stimulation led to an increase in BOLD response in Layer IV of rat barrel cortex, before such an enhancement was observed in any of the other layers, suggesting that layer IV was the first to receive feed forward drive from upstream areas (***Yu et al., 2014***). In contrast, subsequent cortico-cortical connections in the same task appeared to activate Layers II-III and V in the motor cortex and contralateral barrel cortex before this affected any of the other layers, suggesting that these layers were the first to receive feedback signals. To what extent these results generalize to human cortex, however, remains to be investigated.

Recent advancements in fMRI have made it possible to also investigate the functional role of cortical layers in humans (e.g. ***Polimeni et al.*** (***2010***); ***Maass et al.*** (***2014***); ***Kok et al.*** (***2016***); ***Lawrence et al.*** (***2018***); ***Sharoh et al.*** (***2019***)). The human in vivo resolution with fMRI has increased to submillimetre voxel size. The thickness of the cerebral cortex varies between 1 and 4.5 millimetres (***Zilles, 1990***; ***Fischl and Dale, 2000***), giving sufficient resolution to characterise activity across the individual layers. FMRI is now often used to try and measure layer specific activation in, for example, the visual system (***Muckli et al., 2015***; ***Kok et al., 2016***; ***Lawrence et al., 2018***; ***de Hollander et al., 2020***), the motor system (***Huber et al., 2018***), working memory tasks (***Finn et al., 2019***), and to find directional connectivity between language areas (***Sharoh et al., 2019***). If layer specific analysis can make good on its promise of reliably discerning layer specific signals, it can be useful for answering questions in a wide range of cognitive domains (***Lawrence et al., 2019***) and for questions of directional connectivity and cognitive network neuroscience (***Huber et al., 2020***), including research questions involving spatial attention.

While some neurophysiological evidence suggests a differential involvement of the cortical layers in top-down attention (***Nandy et al., 2017***; ***van Kerkoerle et al., 2017***), the effects of attention on the different layers in human visual cortex has remained unclear. Here, we examine with fMRI the potential influence of spatial attention on BOLD activity in the deep, middle and superficial layers in human visual areas V1, V2, and V3. Participants directed their attention to a cued location, and performed an attention-demanding task using an orientation stimulus that was shown at this location, while an unattended grating appeared at a different location of equal eccentricity. On some of the trials, subjects directed their attention to the cued location in anticipation of the stimulus, but no stimulus appeared at this location. We took care to optimise the experimental paradigm, the acquisition, the preprocessing pipeline, the number of subjects, and the statistical analysis to all be state of the art and tailored for an fMRI investigation at laminar resolution. Our expectation was to find deep layer activation in a top-down (attention) condition and middle layer activation in the bottom-up (stimulus) condition, in line with a substantial body of aforementioned histiological, electrophysiological, and fMRI literature. Interestingly, although we observed a reliable increase of the overall BOLD response with attention across all layers, both with and without a stimulus present, we observed no differences in activation level between the layers due to attention. We provide several reasons for these findings in the Discussion.

To facilitate reproducibility of our results, we further include a reproducible and openly accessible processing pipeline for layer-specific analyses (https://doi.org/10.5281/zenodo.3428603 (***Van Mourik et al., 2018***)). The toolbox includes benchmark tests for interactive visualisation of high resolution coregistration, cortical lamination, and cardiac and respiratory noise filtering. To enhance transparency, the data analysis pipeline is furthermore presented as an online visual workflow that can be easily inspected, shared, and adapted to fit any laminar study’s needs. As a result of the relative novelty of fMRI investigations into individual cortical layers, previous work has used a large variety of high-resolution toolboxes and analyis pathways (LAYNII in ***Huber et al.*** (***2017***, 2018); OpenFmriAnalysis in ***Lawrence et al.*** (***2018***); Nighres in ***Huntenburg et al.*** (***2018***); BrainVoyager in ***Goebel*** (***2012***)), which has made direct comparison between studies difficult. We hope our analysis pipeline will help ameliorate this issue, although we readily recognize that other toolboxes and solutions could similarly achieve such goals.

## Results

We first describe the overall task effects that form the basis for our laminar specific fMRI analysis. Our experimental paradigm is depicted in ***Figure 4*** and described in detail in Methods and Materials. In brief, we used an orientation discrimination task in order to investigate the (laminar specific) effects of a visual stimulus and directed spatial attention. Analysis of the behavioural results showed that subjects generally performed well on the task. The mean orientation discrimination threshold across participants was 6.6°.

**Figure 4.**
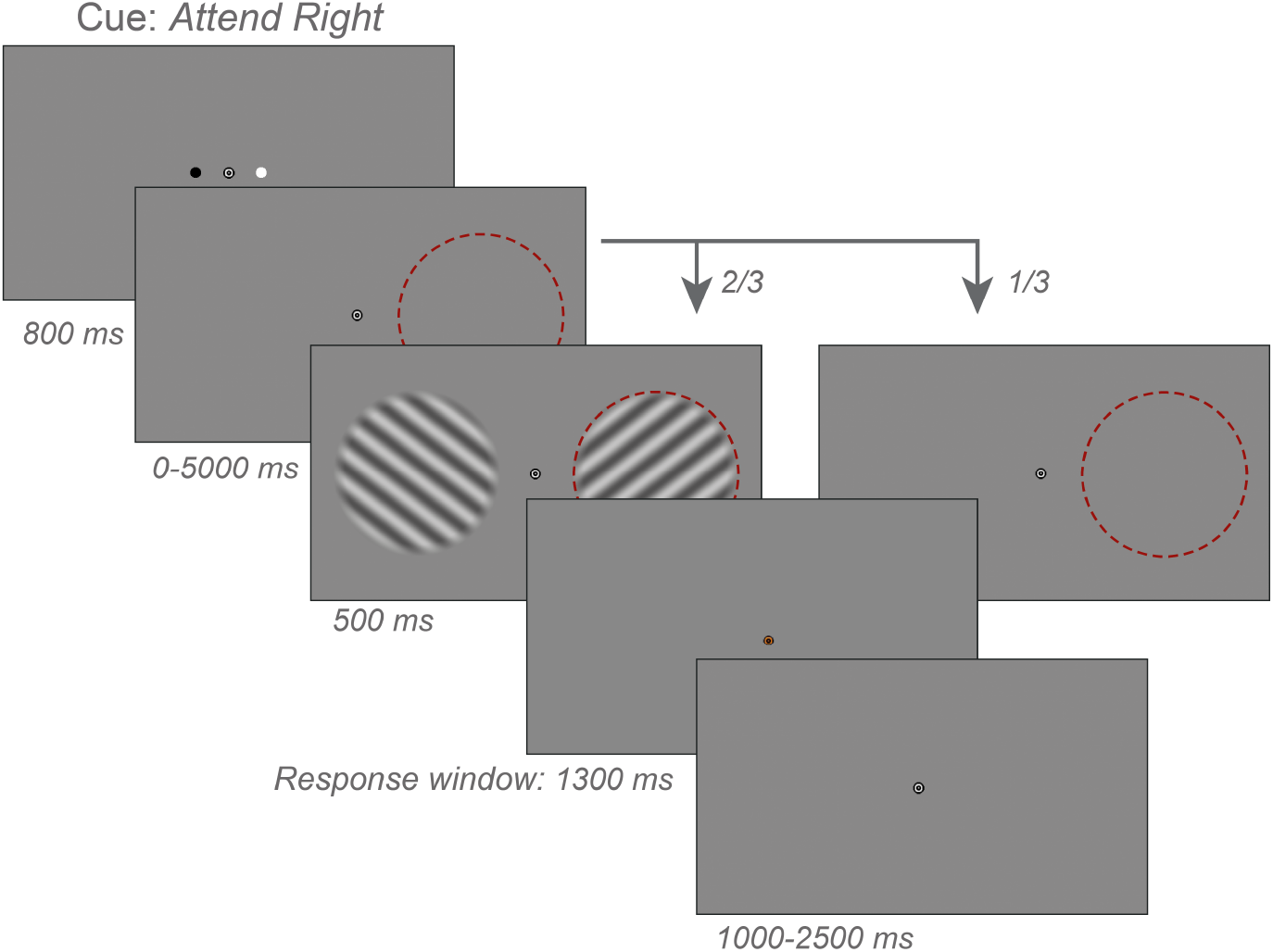
Stimuli and experimental procedure. Example of a trial sequence from the experiment. Subjects fixated a central bull’s eye target while gratings of independent orientation (± 45°) appeared in each hemifield. A compound black/white cue indicated whether subjects should attend to the left or right stimuli; in this example, the white circle indicates ‘attend right.’ Subjects had to discriminate near-threshold changes in orientation of the attended grating with respect to the closest diagonal. In one-third of trials, no stimuli appeared at either location. Red circles depict the attended location and were not present in the actual display.

### Spatial attention increases fMRI response amplitudes

The experiment consisted of four experimental conditions: a two-by-two design in which we manipulated the effect of bottom-up visual stimulation combined with an attentional manipulation across the two hemifields. The stimulus consisted of two orientation gratings presented to the left and right side of fixation. Participants were cued to attend to one location at either side of fixation. They performed a two-alternative-forced-choice task on the stimulus at the attended location, indicating whether its orientation was rotated clockwise or counter clockwise with respect to the nearest diagonal orientation. They maintained fixation on a central bull’s-eye stimulus throughout the experiment. The design is described in detail in the Methods and Materials and in ***Figure 4***.

To benchmark the data, we first determined whether directing attention to a spatial location led to a stronger overall response in the visual cortex. Regions of interest consisted of voxels that were significantly activated by the stimulus in all layers of areas V1, V2, and V3 (see Methods). We compared the amplitude of the BOLD response with and without attention, for trials in which a stimulus was presented and those in which no stimulus appeared on the screen (see ***Figure 3***). Data were analyzed using a General Linear Model (GLM) with area, attention (attended vs. unattended), and stimulus (present vs. absent) as factors (see Methods).

**Figure 3.**
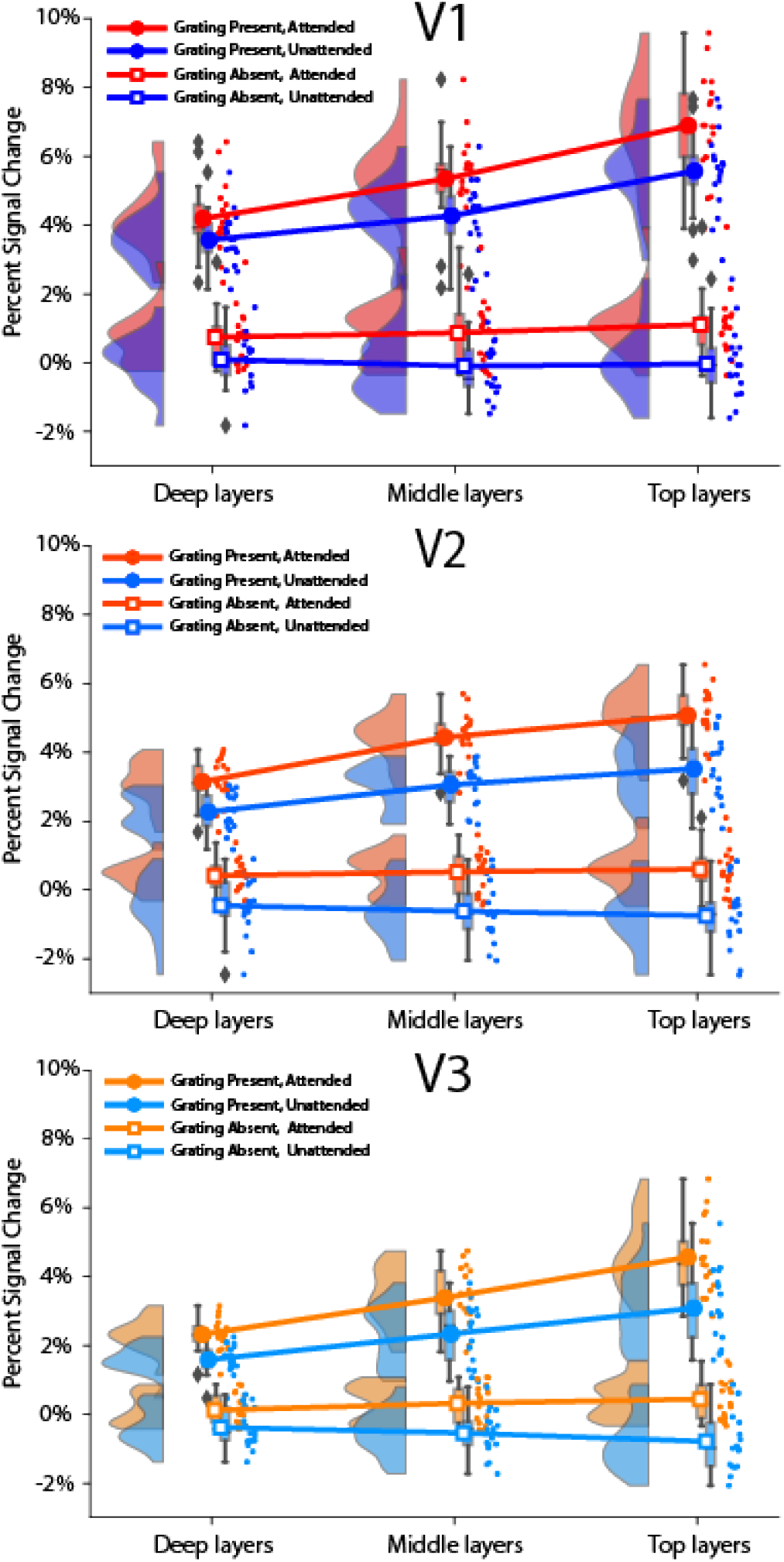
Layer-specific amplitude of the BOLD response in areas V1-V3, for stimuli and locations that were either attended or ignored. Circles indicate when a grating was presented, squares depict when no grating was presented, at either the attended (red) or unattended (blue) location. When a stimulus was presented, activation reliably increased across all layers. Also attention significantly enhanced the BOLD response across all layers.

We first focused on the effects of attention per se. Attention significantly enhanced the BOLD response at the attended location in areas V1-V3 (effect of attention, *F* (1, 16) = 121.3, *p* = 7.07 • 10^−9^). The mean effect sizes (in percent signal change) were 1.01%, 1.09% and 0.96% for V1, V2, and V3 respectively, and slightly stronger to those reported before (e.g., ***Murray*** (***2008***); ***Jehee et al.*** (***2011***); ***Sprague and Serences*** (***2013***)). To account for potential variation in baseline response between visual areas, and facilitate direct comparison with previous studies, we computed an Attentional Modulation Index (AMI; (***Kastner et al., 1999***). The AMI is defined as attentional effects normalised by the summed activity of both attended and unattended conditions. We found AMIs (*μ* ± *σ*) of 0.18 ± 0.05 for V1, 0.24 ± 0.07 for V2, and 0.24 ± 0.09 for V3. Direct comparison between areas revealed that although the absolute contribution of attention did not change, the relative contribution of attention differed significantly between regions (AMI: F(2,32)=10.53, *p* = 3.07 • 10^−4^. Post-hoc comparison showed significant differences in V1 compared to V2: *T* (16) = −4.69, *p* = 2.44 • 10^−3^ and in V1 compared to V3 *T* (16) = −3.17, *p* = 5.89 • 10^−3^, but not in V2 compared to V3: *T* (16) = −0.14, *p* = 0.89). Altogether, these results are in line with previously reported effects of attention on coarse-level BOLD activity in visual cortex (***Somers et al., 1999***; ***Gandhi et al., 1999***), and show that the cortical response for a spatial location is enhanced when attention is directed to that location.

Next, we investigated whether the effects of attention depended on the presence of a visual stimulus. Also in the absence of visual stimulation, there was a significant attention effect (*T* (16) = 9.80, *p* = 3.64•10^−8^), with mean effect sizes of 0.93%. We furthermore observed a slight negative BOLD response in the absence of visual stimulation and when the location was ignored (*T* (16) = −3.12, *p* = 0.0066). This result should be interpreted with caution, however, as the experiment did not include an attention-neutral condition and responses were computed with respect to an implicit baseline response. We next compared attentional effects between trials in which observers were expecting a stimulus but none was presented, and trials in which the stimulus did appear on the screen. We found that the effect of attention in areas V1-V3 was significantly different in the presence compared to absence of visual stimulation (two-way interaction between attention and stimulus, *F* (1, 16) = 6.63, *p* = 0.0204). Specifically, in the presence of a stimulus, the attentional effect was slightly higher (*T* (16) = 2.87, *p* = 0.0060), with no reliable difference between areas (three-way interaction between stimulus, attention and area, *F* (2, 32) = 0.324, *p* = 0.726). Thus, attending to a spatial location clearly enhances the BOLD response at that location, even in the absence of visual stimulation, albeit that attentional effects were sightly reduced when no stimulus was presented to the observer.

To qualitatively assess the shape of the BOLD response over time and confirm the parameters of our GLM approach, we additionally conducted a Finite Impulse Response (FIR) analysis. The FIR analysis can be inspected and reproduced online in a Jupyter Notebook (https://github.com/TimVanMourik/LayerAttention/blob/master/Notebooks/LayerFir.ipynb). We extracted BOLD response curves for each experimental condition (see ***Figure 1***), and observed clear and reliable effects of attention on the BOLD response that were fully consistent with our previous analyses. Because of the left-right modulation of attention, attentional effects were reversed in the left and right hemisphere, as observed before and further illustrated in ***Figure Supplement 1***. Moreover, the cortical response over time for each condition was very well described by the canonical Hemodynamic Response Function (HRF) (*r*^2^ = 0.982 with attention and *r*^2^ = 0.985 without attention). Thus, the HRF model presented a fair description of the cortical responses observed in our experiment.

**Figure 1.**
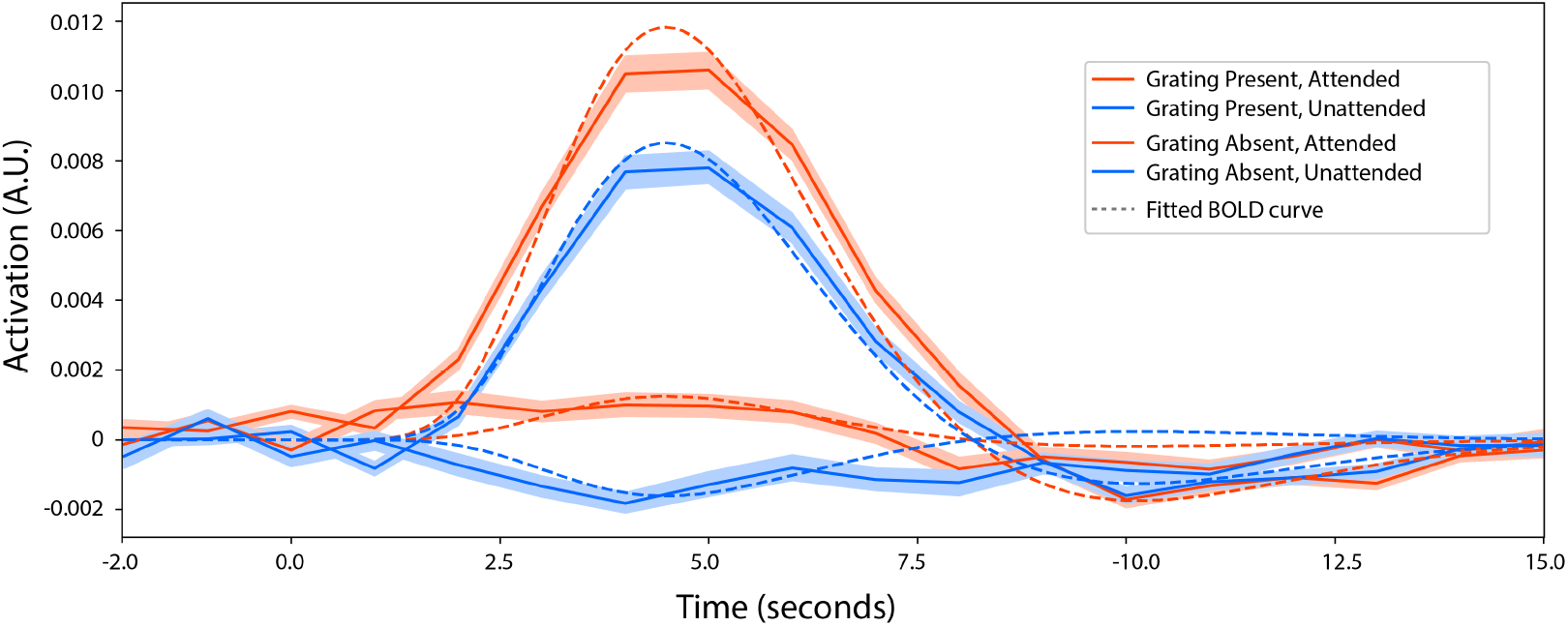
The fitted BOLD response for each experimental condition. The shaded area represents the standard error of the mean over subjects. Results were obtained by fitting a Finite Impulse Response function of 18 time points to the BOLD data, starting at 2 seconds before and running until 15 seconds after stimulus onset. The dashed line indicates an HRF that was fitted to the FIR results of a pilot session. The obtained parameter values were used in the modelled HRF across layers in our GLM analyses (see Methods). These results are collapsed over hemisphere.

### Spatial attention increases responses across the layers

Next, we asked whether attention led to changes in the pattern of activity across cortical layers. For each attention and stimulus condition, we first characterized the BOLD response over time for each of three distinct cortical layers in areas V1-V3 combined. Specifically, we used a FIR analysis to obtain the temporal laminar BOLD response profiles shown in ***Figure 2***. The analysis revealed clear hemodynamic response profiles that were well captured by the canonical HRF. In the presence of a stimulus, there appeared to be a progressive increase in response from the deep to the middle to the top layers, for both the stimulus and attention. In the absence of a stimulus, however, the layer-specific hemodynamic responses did not appear to show noticeable differences.

**Figure 2.**
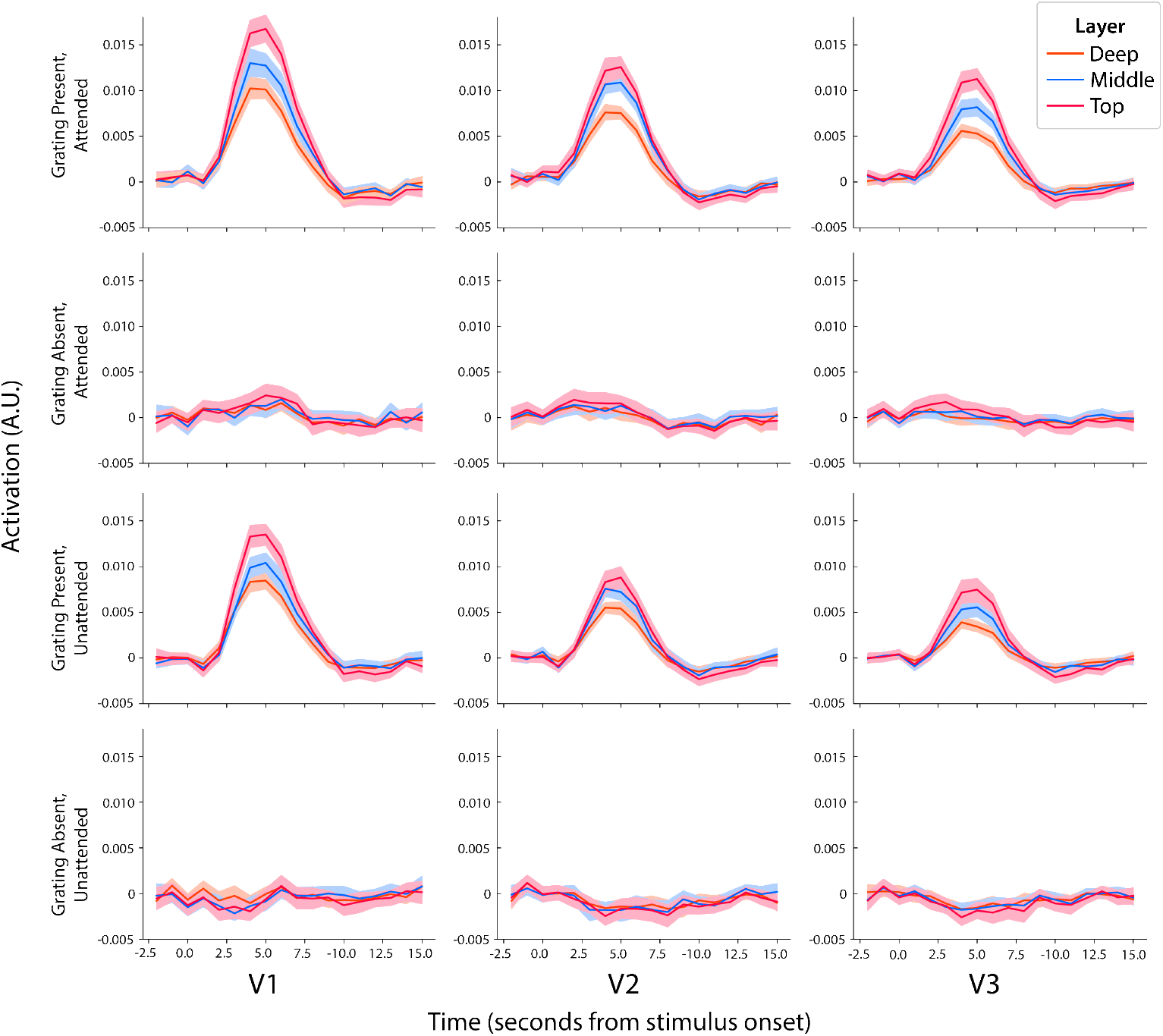
The fitted BOLD response for each experimental condition for three cortical layers. The shaded area represents the standard error of the mean over subjects. The horizontal axis represents time (in seconds) from stimulus onset.

To assess the significance of these effects, data were analyzed using a temporal general linear model with attention, stimulus, area, and layer as factors (see Methods). We found a reliable increase in BOLD response from white matter to pial surface (see ***Figure 3***, overall effect of cortical depth, *F* (2, 32) = 87.5, *p* = 1.07 • 10^−13^). This increase in BOLD response with decreasing distance to the pial surface was reliably larger in the presence of a stimulus (two-way interaction between layer and stimulus, *F* (2, 32) = 85.6, *p* = 1.43 • 10^−13^)). Attention also led to reliable increases in BOLD response with decreasing distance to the pial surface (two-way interaction between layer and attention, *F* (2, 32) = 43.10, *p* = 8.34 • 10^−10^). This increase in BOLD response from lower to higher layers has been observed (e.g. ***Koopmans et al.*** (***2010***); ***Polimeni et al.*** (***2010***); ***Koopmans et al.*** (***2011***); ***Olman et al.*** (***2012***); ***Huber et al.*** (***2018***)) and modeled (***Markuerkiaga et al., 2016***; ***Havlicek and Uludağ, 2020***) previously, and is consistent with the blood flow from the gray-white matter boundary to the pial surface. That is, any change in BOLD response that arises in the deep layers will automatically affect responses in downstream layers, simply because blood flows from deeper to more superficial layers. The accumulation of blood in downstream layers can result in a larger slope of BOLD activation across the layers, even when there is no change in neural activity in these layers.

Next, we determined whether the layer-specific increase in BOLD signal was different for attention-based effects compared to those of the visual stimulus. If so, then this would be consistent with a targeted effect of attention on one of the layers. Interestingly, the attention-based increase in activity was reliably different from stimulus-driven changes in layer response (post hoc comparison between layer by stimulus effect and layer by attention effect; *T* (16) = 9.28, p=7.64 • 10^−8^). This could potentially reflect a specific effect of attention on one of the layers. However, there is also an alternative interpretation, as the magnitude of an interaction effect is tightly coupled to the strength of its main effects in layer-specific analyses. This is because the interaction effect can be interpreted as a difference in slope (signal by depths) between two lines. Due to cortical signal leakage as a result of physiological (blood flow) and methodological reasons (errors in depth measurement), a higher visual stimulus response than an attentional response will be visible in *both* the main effect *and* the cortical slope of the response (see also above). Any reliable difference in slope is picked up as a significant difference between layer by stimulus effect and layer by attention effect. However, this difference in slope would also arise if there was no layer-specific change in neural activity, so this finding alone does not indicate conclusive layer specific activation. Such conclusions might be made if a reliable three-way difference between the effects of attention and stimulus between any of the three layers would be present. However, we did not find a significant three-way interaction (three-way interaction between layer, stimulus and attention, *F* (2, 32) = 0.96, *p* = 0.393), making it difficult to draw any firm conclusions regarding the layer-specific effects of the stimulus versus attention.

While we observed no clear and unambiguous laminar differences across visual areas, could there be a change within a given area? When analyzing stimulus activity, the pattern of activity across the layers was, indeed, significantly different between the three areas (three-way interaction between layer, stimulus, area: F(4, 64)=3.10, p=0.021)). Post hoc analyses revealed that the stimulus response across layers was slightly steeper in V1 than V2 (stimulus by layer effect in area V1 compared to V2, *T* (16) = 2.26, *p* = 0.038), while no significant difference was observed for area V1 compared to area V3 (*T* (16) = 1.35, *p* = 0.196), or V2 compared to V3 (*T* (16) = −1.43, *p* = 0.172). This might reflect a slightly higher top layer activation in V1 compared to V2. On the other hand, it could also reflect the stronger main effect of stimulus in V1, leading to an increased slope of activation over layers, in line with expectations based on blood flow. We then focused on the layer-specific effects of attention in each individual area. Attention does not vary layer specifically per region (attention by layer by region effect *F* (64, 4) = 0.998, *p* = 0.416).

Thus, while the overall effects on BOLD activity of both visual stimuli and attention were robust and similar to previously reported values for visual cortex (e.g. (***Kastner et al., 1999***; ***Jehee et al., 2011***; ***Koopmans et al., 2010***)), both across and within layers, it proved more difficult to interpret the layer-specific pattern of activity for the two conditions. Although at first sight, it may appear that the stimulus affected the layers in a manner different from attention (when analyzed across areas), the effect should be interpreted with caution because the stimulus also led to a stronger coarse-level cortical response than attention. We wanted to rule out that some important parameter choices obscured true effects. We verified robustness of our main results by changing the size of the region of interest and by varying the most important parameters of our layer extraction technique. We reprocessed the data and recomputed our layer and region specific statistical analysis for the experimental conditions. The control analyses established that the results were not strongly affected by the number of voxels included in the analyses (***Figure Supplement 1*** and ***Figure Supplement 2***), nor by the number of layers analyzed (***Figure Supplement 3***). In addition, the results did not qualitatively change when layer activation profiles were defined using volume interpolation (***Figure Supplement 4***) as opposed to using a laminar spatial GLM as we did in the main analyses (see Methods). The control analyses and their comparison with the main analysis is further described in the supplemental materials.

All figures and reported statistics are available as Jupyter Notebooks (https://doi.org/10.5281/zenodo.3428603). The (fully anonimised) BOLD time courses are included such that all results can be readily reproduced.

## Discussion

This study investigated the effects of spatial attention on the BOLD signal measured from individual layers in early visual cortex. Focusing first on the overall amplitude of the BOLD response in all layers combined, we found that attending to a stimulus reliably and substantially increased the BOLD signal in early visual areas, both when a stimulus was presented to the observer and in the absence of physical stimulation (cf. (***Kastner et al., 1999***; ***Murray, 2008***; ***Li et al., 2008***)). Moreover, and much in line with earlier results on layer-specific activation patterns in visual cortex (***Koopmans et al., 2010***; ***Polimeni et al., 2010***), we observed a general increase in activation towards the superficial layers, which is likely caused by greater susceptibility to draining veins on the pial surface (***Koopmans et al., 2011***). Interestingly, and much to our surprise, we observed no forthright differential activity in the individual layers when comparing between top-down (attention-driven) and bottom-up (stimulus-driven) activity - a finding that stands in notable contrast to some previous observations. We discuss several potential reasons for the discrepancy in findings below, and suggest a more standardized approach to laminar fMRI analysis as a potential solution that could boost agreement in research findings.

Why did we not find a targeted effect of attention on one of the layers? One possibility is that our data are simply insufficiently robust to show a significant difference in activity across depth between the two conditions. It is well known that the BOLD signal includes multiple sources of noise related to both MRI scanner and participant, and this holds especially true for signals recorded at the sub-millimeter scale. For example, at a resolution this high, even the smallest movement of the participant may cause additional blurring of the data, with potentially detrimental effects on the signal-to-noise ratio. For this reason, we collected data from 17 participants - a sample size much larger than typical in attention-based fMRI studies at standard spatial resolution (cf. N=4-6 in ***Kastner et al.*** (***1999***); ***Kamitani and Tong*** (***2005***); ***Jehee et al.*** (***2011***)), and on the high end compared to layer-based fMRI studies at high resolution (cf. N=6 in ***Polimeni et al.*** (***2010***), N=4 in ***Muckli et al.*** (***2015***), N=10 in ***Kok et al.*** (***2016***), N=12 in (***Klein et al., 2018***), N=21 in (***Lawrence et al., 2018***), N=22 in (***Sharoh et al., 2019***)). To minimize the effects of various sources of noise, we took great care in measuring and removing physiological artifacts, and further improved existing layer extraction techniques by developing a novel technique to separate laminar signal from different layers (***Van Mourik et al., 2019***). Indeed, the combined success of these procedures is well illustrated by the effect sizes observed in the current study for both stimulus presentation (4.5%, 3.3%, 2.8% in V1, V2 and V3) and attention (0.41%, 0.64%, 0.59% in V1, V2 and V3), which are comparable or higher to those reported in previous work (***Murray, 2008***; ***Jehee et al., 2011***). We also ensured that similar results were obtained using more conventional layer-extraction procedures, and that the results were robust to changes in signal extraction method or number of layers. There are, however, some differences in experimental design between our study and previous laminar investigations that could potentially account for the incongruity in results. Because we were interested in the degree to which top-down processes could be dissociated from feed forward stimulation with fMRI, we directly contrasted between these two conditions in our analyses. Previous studies, on the other hand, have focused on top-down activity in isolation (e.g. ***Muckli et al.*** (***2015***), ***Kok et al.*** (***2016***), ***Lawrence et al.***(***2018***)). Two-by-two experimental designs are surprisingly rare in layer-specific analysis and have only been recently employed by ***de Hollander et al.*** (***2020***). We emphasise that a multifactorial design is important to account for changes due to, for example, cortical depth and signal leakage per se, as opposed to true layer-based changes in activity due to the experimental manipulations. Alternate strategies are to compare between layers and conditions in terms of information content (***Muckli et al., 2015***), retinotopic preference (***Klein et al., 2018***), or by focusing on inter-regional laminar communication (***Sharoh et al., 2019***). For example, ***Klein et al.*** (***2018***) recently observed a cortical depth-dependent shift in population receptive fields with spatial attention. This raises the intriguing possibility that, in our study, spatial attention led to a depth-dependent shift in the strength of relatively fine-grained orientation-selective responses – indeed, previous coarse-scale fMRI studies have observed that orientation selectivity can change even when there is no change in amplitude across the population (***Jehee et al., 2011***, ***2012***).

We initially hypothesised that attention provides additional information about the stimulus (e.g. knowledge about its location). This information would come from higher level areas, and would presumably affect the deep or superficial layers. This was suggested by previous work using high-resolution fMRI focussing not on spatial attention, but rather figure-ground segmentation (***Kok et al., 2016***), and other extra-classical receptive field effects in cortex (***Muckli et al., 2015***). However, our results are incongruent (insofar that they are comparable) and do not find similar effects for spatial attention. It is conceivable, however, that processes of perceptual grouping operate on the individual cortical layers in a manner different from the spatial attentional mechanisms studied here. It is known from primate studies, for example, that attention increases the response gain of neurons in visual cortex (***Treue and Trujillo, 1999***; ***Martinez-Trujillo and Treue, 2004a***) - such an increase in attentional gain could lead to general enhancements in neural activity irrespective of cortical layer, as we have observed here.

Regardless of the potential reasons for the disparity between current and previous results, we believe our study presents an important message to a field that is currently in its nascent stages of development. We hope that the results and procedures detailed here will help move the field forward and resolve which experimental parameters are paramount to detecting differential activity between individual layers in human visual cortex with high-resolution fMRI. To facilitate comparison between results, and provide loose analysis guidelines for future laminar studies, we have furthermore provided all our analysis code and data online (see Data Avalibility section), in a format that is visually insightful and analytically meaningful. We hope that our software and analysis pathways will prove to be a useful resource, and boost the comparability and replicability of future results.

## Methods and Materials

### Participants

Nineteen healthy adults (aged 22-27, eight female), with normal or corrected-to-normal vision, participated in this study. All participants provided written informed consent in accordance with the guidelines of the local ethics committee (CMO region Arnhem-Nijmegen, the Netherlands, and ethics committee of the University Duisburg-Essen, Germany). Two subjects were excluded from analysis; one subject was excluded due to insufficient performance on the orientation discrimination task (their behavioral performance was at chance-level), and another due to weak retinotopic maps. The remaining data from 17 subjects were analyzed.

### Experimental design and stimuli

Observers viewed the visual display through a mirror mounted on the head coil. Visual stimuli were generated by a Macbook Pro computer running MATLAB and the Psychophysics Toolbox software (***Brainard, 1997***; ***Pelli, 1997***), and displayed on a rear-projection screen using a luminance-calibrated EIKI projector (resolution 1,024 × 768 pixels, refresh rate 60 Hz). Participants were required to maintain fixation on a central bull’s-eye target (radius: 0.25°) throughout each experimental run. Each run consisted of an initial fixation period (3000 ms) followed by 32 stimulus trials (average trial duration: 4.7 seconds). Trials were separated by inter-trial intervals of variable duration (1000-2500 ms, uniformly distributed across trials). Each trial started with the presentation of a central attention cue (800 ms). This was followed by a delay period of variable duration (0-5000 ms; drawn from an exponential distribution to ensure a constant hazard rate), after which the two orientation stimuli appeared on the screen (500 ms; two-thirds of trials). The orientation stimuli were followed by a response window (1300 ms), in which the fixation target turned orange, and observers indicated their response by pressing a button with their right index or middle finger. On one-third of the trials, no orientation stimuli appeared and the screen remained blank for the remainder of the trial. A single trial of the experiment is illustrated in ***Figure 4***.

Stimuli were two counterphasing sinusoidal gratings of independent orientation ~45°or ~135°; size: 7°; spatial frequency: 1 cycle per °; randomized spatial phase; contrast: 50%; contrast decreased linearly to 0 towards the edge of the stimulus over the last degree), centered at 5°to the left and right of fixation. We used a compound white/black cue consisting of two dots (dot size 0.25°) that straddled the fixation point (0.8°to the left and right of fixation) to indicate with 100% validity which of the two gratings should be attended (***Jehee et al., 2011***). Subjects were instructed to attend to the same side of fixation as either the white or black dot in the compound cue. Participants were instructed to detect a small clockwise or counterclockwise rotation in the orientation of the grating at the attended location with respect to a base orientation at 45°or 135°. The size of rotation offset was adjusted with an adaptive staircase procedure using QUEST (***Watson and Pelli, 1983***), such that participants detected approximately 80% of the offsets correctly.

All but one participant completed 18 stimulus runs. The remaining participant completed 12 runs due to equipment failure. Retinotopic maps of visual cortex were acquired in a separate scan session at a 3T scanner using conventional retinotopic mapping procedures (***Sereno et al., 1995***; ***DeYoe et al., 1996***; ***Engel et al., 1997***).

### MR data acquisition

Functional images were acquired on a Magnetom Siemens 7T scanner with a 32-channel head coil (Nova Medical, Wilmington, USA) combined with dielectric pads (***Teeuwisse et al., 2012***), using a 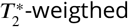 3D gradient-echo EPI sequence (***Poser et al., 2010***) (TR/TE/*α*=3060 ms/20 ms/14°, 72 slices oriented orthogonally to the calcarine sulcus, voxel size [0.8 mm]3, FOV: [192 mm]^2^, GRAPPA factor 8).

Gradient maximum amplitude was 40 mT/m (in practice, however, this maximum was not reached) and the maximum slew rate was 200 T/m/s. Shimming was performed using the standard Siemens shimming procedure for 7T. There were 18 runs of 72 ± 4 volumes. As the lengths of the events and the inter trial interval were of unequal length, there was a small variation in the number of volumes per run.

Finger pulse was recorded using a pulse oximeter affixed to the index finger of the left hand. Respiration was measured using a respiration belt placed around the participant’s abdomen.

Anatomical images were acquired using an MP2RAGE sequence (***Marques et al., 2010***) [0.75 mm]^3^, yielding two inversion contrasts (TR/TE/TI1/TI2 = 5000 ms/1.89 ms/900 ms/3200 m).

In a separate session prior to the main experiment, a retinotopy session was conducted at a Siemens 3T Magnetom Trio scanner. A high-resolution T1-weighted anatomical scan was acquired (MPRAGE, FOV 256 × 256, 1 mm isotropic voxels) at the start of the session. Functional images were subsequently collected using 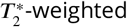 gradient echo EPI, in 30 slices oriented perpendicular to the calcarine sulcus (TR/TE/*α* = 2000 ms/30 ms/90°, FOV = 64 × 64, [2.2 mm]3 isotropic resolution).

### Functional MRI preprocessing

#### Data preprocessing

Data were corrected for subject motion using SPM with the mean functional volume across time as a reference (***Friston et al., 1995***). Residual motion-induced fluctuations in the BOLD signal were removed through linear regression, based on the alignment parameters (3 translation and 3 rotation parameters, no derivatives) of SPM. Scanner drifts were corrected via linear regression with high-pass filter regressors to filter out frequencies below 1/64 Hz. Pulsating signals as a result of the respiratory and cardiac cycle were removed as follows. The cardiac/respiratory peaks were automatically detected from the physiological recordings using in-house interactive peak-detection software, and manually corrected where needed. With a custom MATLAB implementation of RETROICOR (***Glover et al., 2000***), fifth order Fourier regressors were constructed for heart rate and respiration and subsequently removed from the functional images via linear regression. A small part (10% of respiratory measurements, and 18% of heart rate measurements) was of insufficient quality and could not be used in this analysis. Functional data for these time frames were used in the main analysis but uncorrected for cardiac and respiratory noise.

The functional and anatomical scans were brought to the same space by registering the anatomical surface from the retinotopy session to the mean functional volume using boundary based registration (BBR), implemented in FreeSurfer’s bbregister (***Greve and Fischl, 2009***). All registration results were inspected and manually refined when necessary. Where needed, registration was improved by an additional pass of BBR using an in-house MATLAB implementation. Local distortions in EPI due to field inhomogeneity were corrected by means of recursive boundary registration (***Van Mourik et al., 2019***), which recursively applies BBR to small portions of the cortical surface to correct topology locally by means of optimizing the grey-white matter contrast along the surface. We used 7 layers of recursion and only looked for translations and scalings in the phase encoding direction. Note that this procedure displaces the *surface mesh* and not the volume, so this has no smoothing effect on the (layer) signal.

Because of temporal changes in magnetic field inhomogeneity, local topology slightly changed over the course of the entire session. For this reason, the 18 functional runs obtained for each subject were first divided into three groups of each 6 contiguous runs, and then each group was pre-processed separately. Time courses were subsequently concatenated before entering the main analyses.

#### Regions of Interest

Regions of interest (areas V1, V2, V3) were defined on the reconstructed cortical surface using standard retinotopic mapping procedures (***Sereno et al., 1995***; ***DeYoe et al., 1996***; ***Engel et al., 1997***). After identifying areas V1-V3, data from the main experiment were smoothed along the reconstructed cortical surface with a Gaussian kernel (FWHM: 4 mm). The smoothed version of the data was only used in region of interest selection, and not in the main analysis. In each area, we then selected the 600 vertices that responded most strongly to the stimulus (shown on the cortical surface in ***Figure Supplement 1***). The selected vertices were resampled from the cortical surface back to subject space by means of FreeSurfer’s label2vol. T-values of selected voxels (*μ* ± *er*) were V1: T=2,989 ± 0.854, V2: T=2.317 ± 0.689 and V3: T=2.117 ± 0.713). Note that the selection of voxels based on visual activation per se is orthogonal to the analysis of interest, which addresses the effects of attention on individual layers in cortex. Control analyses verified that our results were not strongly affected by the number of vertices selected for subsequent analysis (See ***Figure Supplement 1***).

#### Cortical profile extraction

Layer specific signals were obtained by means of a layer specific spatial General Linear Model (GLM) as described in detail and proposed by (***Van Mourik et al., 2018***). In brief, we obtained three equivolume layers, following the procedures described in (***Waehnert et al., 2014***). We took the reconstructed cortical surface as determined by FreeSurfer (***Dale et al., 1999***) as a basis for this analysis. We used a custom implementation of ***Waehnert et al.*** (***2014***) with mild adaptations: the gradient and the curvature of the cortex were defined as a function of Laplacian streamlines in the grey matter as this more naturally follows the structure of cortical columns (***Leprince et al., 2015***). Partial volume inaccuracies were adjusted for by explicitly taking into account the orientation of the voxel with respect to the cortex (***Van Mourik et al., 2019***).

The procedure enabled us to divide the gray matter in three equivolume cortical layers, which amounts to roughly one voxel per layer. We additionally defined a volume on either side of these three cortical layers to capture signals for white matter and cerebrospinal fluid. On the basis of these definitions, we then computed a laminar mixing matrix of layer signal over the voxels (***Van Mourik et al., 2018***). This was used as a *spatial* design matrix to unmix the layer signal. By means of a spatial regression of this matrix against the functional data within the ROIs, we obtained laminar time courses.

In separate control analyses, we verified that our laminar results did not qualitatively depend on the specific methods that were used to extract the laminar activation profile. We varied several parameters: the number of layers that we extracted and the method of obtaining laminar signal. For the former, we computed the cortical layers with the spatial GLM for four layers instead of three. For the latter, we computed the laminar signal based on a more conventional interpolation approach: by means of interpolating the fMRI volumes at three points in between the cortical surfaces we obtained laminar signal. This may contain contamination from other layers, but is impervious to potential estimation errors as a result of the laminar spatial regression. The difference between approaches is described in detail in ***Van Mourik et al.*** (***2018***).

### Temporal analysis

Temporal linear regression was used to compare between the experimental conditions. Regressors were created as follows. The stimuli appeared during the stimulus window on 2/3rds of trials, which were modeled with a single regressor (stimulus on). The remaining stimulus windows were also modeled with a regressor (stimulus off). In addition, attention could either be directed to the left or right visual field; these conditions were each modeled with a regressor. We so obtained four regressors for each of the conditions of interest. Thus, for a given retinotopic region of interest, the four different conditions were: stimulus, no stimulus, attended, and unattended. We used a double-gamma function, as defined by SPM (parameters: time-to-peak first gamma: 5 second, time-to-peak second gamma: 10 seconds, amplitude ratio: 2:1), to model the fMRI responses. These parameters were established based on an initial finite impulse response (FIR) analysis (***Friston et al., 1998***) using the visual stimulus response from four pilot subjects (not included in the current study). Based on the observed fMRI response in this pilot data set, temporal or dispersion derivatives were not included into the statistical model of the main experiment. The baseline signal of each run was captured by adding a regressor column of ones for each run separately. As described above, the data were pre-processed by means of nuisance regression. This was performed by adding the nuisance regressors to the design matrix, effectively adjusting for the statistical loss in degrees of freedom as a result of nuisance regression. The reference of one percent signal change was the height of a peak of a two-second-long isolated event (***Mumford, 2007***).

We additionally performed a Finite Impulse Response (FIR) analysis to qualitatively assess the BOLD response over time for each of the four conditions. This was used to confirm that the used HRF would accurately describe the true BOLD response. To visualize the cortical response over time for each of the four conditions, we analyzed the data using FIR filters (***Friston et al., 1998***), applied to each layer. Specifically, we constructed FIR regressors for each of the four experimental conditions, each containing 18 time points that represented a time window of 1 second, starting 2 seconds before stimulus onset and running until 15 seconds thereafter.

We started at the level of the cortical region, initially with no further specification into layers. The temporal regressions were performed on the previously extracted time course of V1, V2, and V3. The obtained parameter estimates were divided by their baseline estimates, in order to convert them to percent signal change. The values in percent signal were compared at the group level by means of ANOVAs and t-tests as appropriate. As the experiment was left-right symmetric and we found no differences between hemispheres in the analyses of interest, the hemispheres were treated as two measurements per participant.

We subsequently focused on the laminar level. For a qualitative assessment of the layer specific BOLD response, we repeated the FIR analysis for each experimental condition and each layer. These qualitative results are shown in ***Figure 2***. The BOLD responses do not seem to vary per layer beyond a general intensity increase towards the pial surface. To further investigate this quantitatively, we repeated the region specific analysis with the addition of the ‘layer’ factor. By means of an ANOVA, we ascertained layer specific effects and their interactions with the stimulus and the region effects. These were followed by t-tests where appropriate, to further inspect significant results. As the experiment was left-right symmetric and we found no differences between hemispheres in the analyses of interest, we took the hemisphere data as two measurements per participant.

## Code and data availability

All code and data can be found online: https://doi.org/10.34973/bf42-rx14 for a full data set for a single subject and all raw files from the scanner https://doi.org/10.34973/eb4d-md15. Layer-specific analysis were performed using custom-written software available online https://github.com/TimVanMourik/OpenFmriAnalysis. This pipeline can be inspected graphically (***Van Mourik et al., 2018***) (https://giraffe.tools/workflow/TimVanMourik/LayerAttention). Moreover, it can readily be applied to custom data; we prepared data from a representative subject to be used as a template pipeline.

Preprocessing of the data and construction of design matrices was performed in MATLAB. The FIR analysis, the region of interest analysis, and the layer specific analyses were performed in an openly available Jupyter notebook (available at https://doi.org/10.5281/zenodo.3428603).

## Author Contributions

PJK and JFMJ designed the experiment. TvM and LJB collected the data. TvM and DGN developed the laminar analysis techniques. TvM and JFMJ analysed the data. TvM, DGN, and JFMJ wrote the paper.

## Appendix 1 Layer specific HRF for all conditions and layers

To assess the degree to which the results are robust to different analysis choices, we additionally analysed the data using a variety of alternative methodological parameters. This section discusses the obtained results. Our main analysis was done on a region of interest of 600 vertices. The method of extracting cortical signal was the spatial GLM (***Van Mourik et al., 2019***) with three cortical layers. We redid the main analysis with a smaller region of interest (300 vertices) and a larger one (900 vertices), other factors remaining equal. We further wanted to make sure that changing the number of layers did not qualitatively change our interpretation and recomputed the main analysis with four instead of three cortical layers. For comparison with traditional studies that use signal interpolation for obtaining laminar signal, we also employed this technique in an additional control analysis. All results (p-values) of the main analysis, the ANOVA of the factors stimulus, attention, layer, and region, are included in one table ( 1).

For further inspection, the results are also included as figures, analogous to ***Figure 3*** in the main text. ***Figure Supplement 1*** and ***Figure Supplement 2*** show stimulus and attention-based effects across layers after selecting, respectively, the 300 and 900 most activated vertices (cf. ***Figure 3*** in the main text, based on 600 vertices). Note that, per this selection, these results should (and do) show higher and lower activation values for the top 300 and 900 vertices, respectively. ***Figure Supplement 3*** shows results after defining four cortical layers, rather than three, and Figure ***Figure Supplement 4*** depicts results obtained from interpolation instead of a laminar spatial GLM. Error bars indicate ±1 SEM. In all Figures, presenting a stimulus (circles) resulted in a reliable increase in BOLD response from deep to superficial layers. The BOLD response was significantly enhanced for the attended (red) compared to unattended location (blue) across layers, both when a stimulus was presented, and in the absence of visual stimulation (see Table 1 for statistics).

We repeated the main analysis four additional times. Thus, it is to be expected that some p-values that are around the significance threshold in the main analysis, fall slightly below or rise slightly above it in some of the control analyses. There were only two instances where significance (*p* < 0.05) changed compared to the layer analyses that are presented in the main text.

While we observed a trending effect for the stimulus by attention interaction in the main laminar analysis (trending, with *p* = 0.0733), it was significant for the analysis with a larger ROI (*p* = 0.0272) and with four extracted layers (*p* = 0.0118), but not for the smaller ROI (*p* = 0.271) and trending for the signal extraction by means of interpolation (*p* = 0.0552). These results are all close together and hovering around the significance threshold of *p* = 0.05. A stimulus by attention effect ought to be interpreted as a multiplicative effect of stimulus and attention, i.e. an additional signal increase when the presented stimulus is attended compared to when the location per se (i.e. no stimulus) is attended. However, this trending effect does not affect any of our conclusions regarding the layer-specific effects of stimulus and attention. While the Stimulus by Layer by Area interaction was significant in the main analysis (p = 0.0214), this interaction was not significant after selecting the 300 most activated vertices (p = 0.2056). However, in the three other control analyses, the effect became more pronounced (larger ROI: *p* = 8.89 • 10^−3^, interpolation: *p* = 2.78 • 10^−5^, and four layers: *p* = 6.19 • −4).

**Appendix 1 Table 1.**
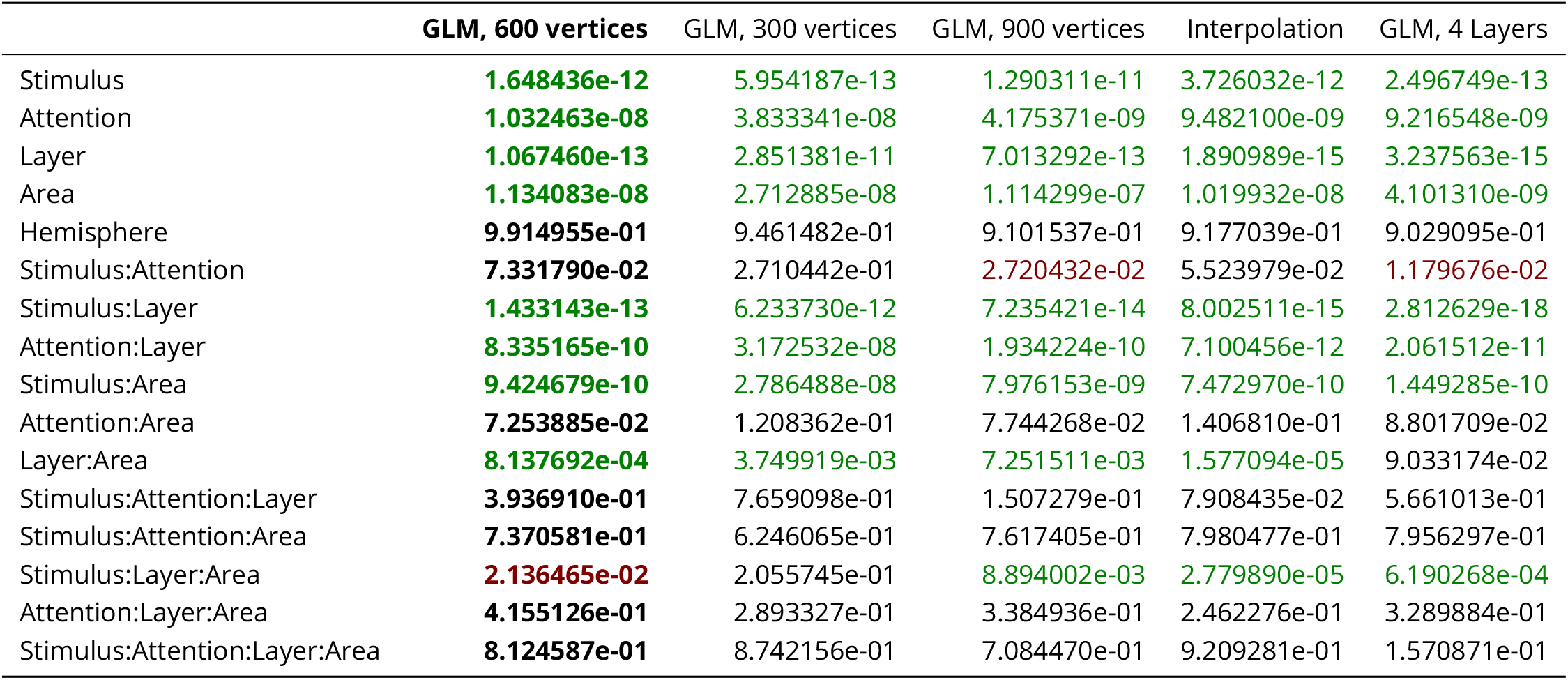
The p-values for the ANOVA as described in the body of the paper. The five columns are the main analysis (first column, bold face), and four control analyses: an analysis based on a smaller ROI (300 vertices); on a larger ROI (900 vertices); the same ROI but laminar signal extracted by means of interpolation instead of a GLM; and a GLM but with four layers instead of three. P-values above 0.05 are marked in black. P-values between 0.05 and 0.01 are marked in red. P-values below 0.01 are marked in green.

|  | **GLM, 600 vertices** | GLM, 300 vertices | GLM, 900 vertices | Interpolation | GLM, 4 Layers |
| --- | --- | --- | --- | --- | --- |
| Stimulus | **1.648436e-12** | 5.954187e-13 | 1.290311e-11 | 3.726032e-12 | 2.496749e-13 |
| Attention | **1.032463e-08** | 3.833341e-08 | 4.175371e-09 | 9.482100e-09 | 9.216548e-09 |
| Layer | **1.067460e-13** | 2.851381e-11 | 7.013292e-13 | 1.890989e-15 | 3.237563e-15 |
| Area | **1.134083e-08** | 2.712885e-08 | 1.114299e-07 | 1.019932e-08 | 4.101310e-09 |
| Hemisphere | **9.914955e-01** | 9.461482e-01 | 9.101537e-01 | 9.177039e-01 | 9.029095e-01 |
| Stimulus:Attention | **7.331790e-02** | 2.710442e-01 | **2.720432e-02** | 5.523979e-02 | **1.179676e-02** |
| Stimulus:Layer | **1.433143e-13** | 6.233730e-12 | 7.235421e-14 | 8.002511e-15 | 2.812629e-18 |
| Attention:Layer | **8.335165e-10** | 3.172532e-08 | 1.934224e-10 | 7.100456e-12 | 2.061512e-11 |
| Stimulus:Area | **9.424679e-10** | 2.786488e-08 | 7.976153e-09 | 7.472970e-10 | 1.449285e-10 |
| Attention:Area | **7.253885e-02** | 1.208362e-01 | 7.744268e-02 | 1.406810e-01 | 8.801709e-02 |
| Layer:Area | **8.137692e-04** | 3.749919e-03 | 7.251511e-03 | 1.577094e-05 | 9.033174e-02 |
| Stimulus:Attention:Layer | **3.936910e-01** | 7.659098e-01 | 1.507279e-01 | 7.908435e-02 | 5.661013e-01 |
| Stimulus:Attention:Area | **7.370581e-01** | 6.246065e-01 | 7.617405e-01 | 7.980477e-01 | 7.956297e-01 |
| Stimulus:Layer:Area | **2.136465e-02** | 2.055745e-01 | 8.894002e-03 | 2.779890e-05 | 6.190268e-04 |
| Attention:Layer:Area | **4.155126e-01** | 2.893327e-01 | 3.384936e-01 | 2.462276e-01 | 3.289884e-01 |
| Stimulus:Attention:Layer:Area | **8.124587e-01** | 8.742156e-01 | 7.084470e-01 | 9.209281e-01 | 1.570871e-01 |

---

Figure 1-Figure supplement 1. The results for the left and right hemisphere separately are displayed in the supplementary figures

**Figure 1-Figure supplement 1.**
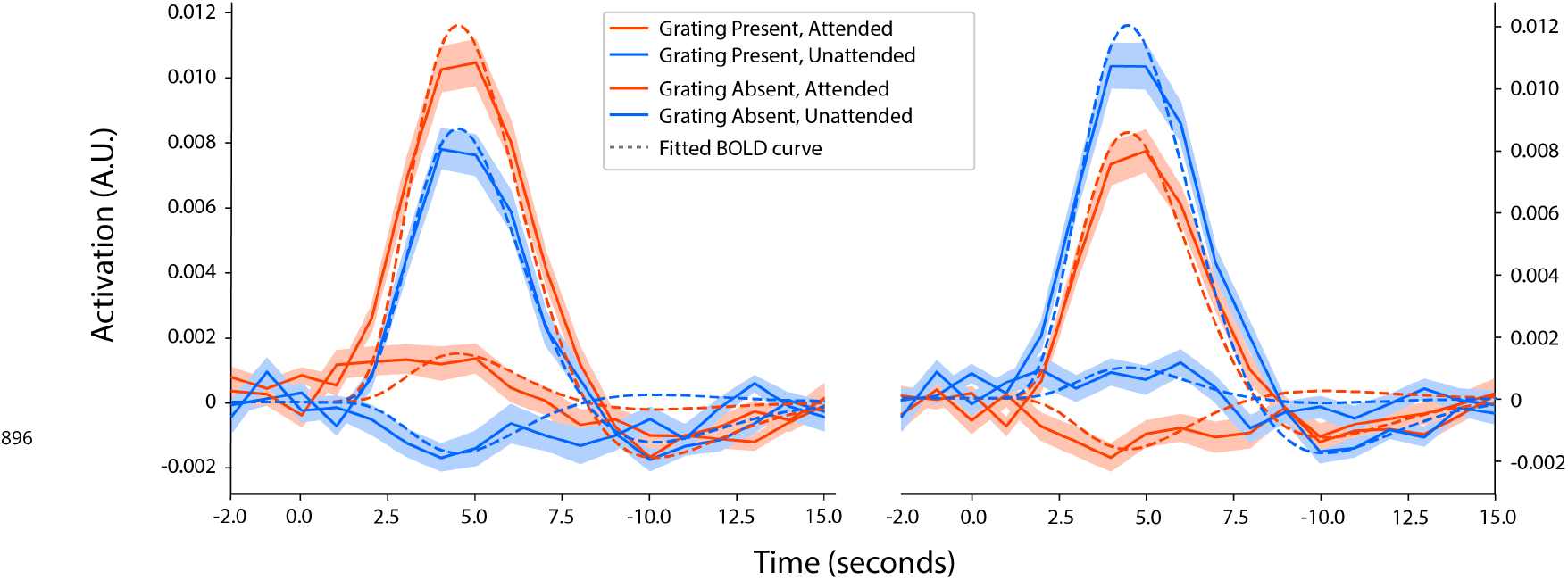
The fitted BOLD response for each experimental condition, separated by hemisphere. The shaded area represents the standard error of the mean over subjects. Results were obtained by fitting a Finite Impulse Response function of 18 time points, starting at 2 seconds before and running until 15 seconds after stimulus onset. The dashed line indicates an HRF that was fitted to the responses in a pilot session (see Methods). The same HRF parameter values were used in other statistical analysis.

**Figure 3-Figure supplement 1.**
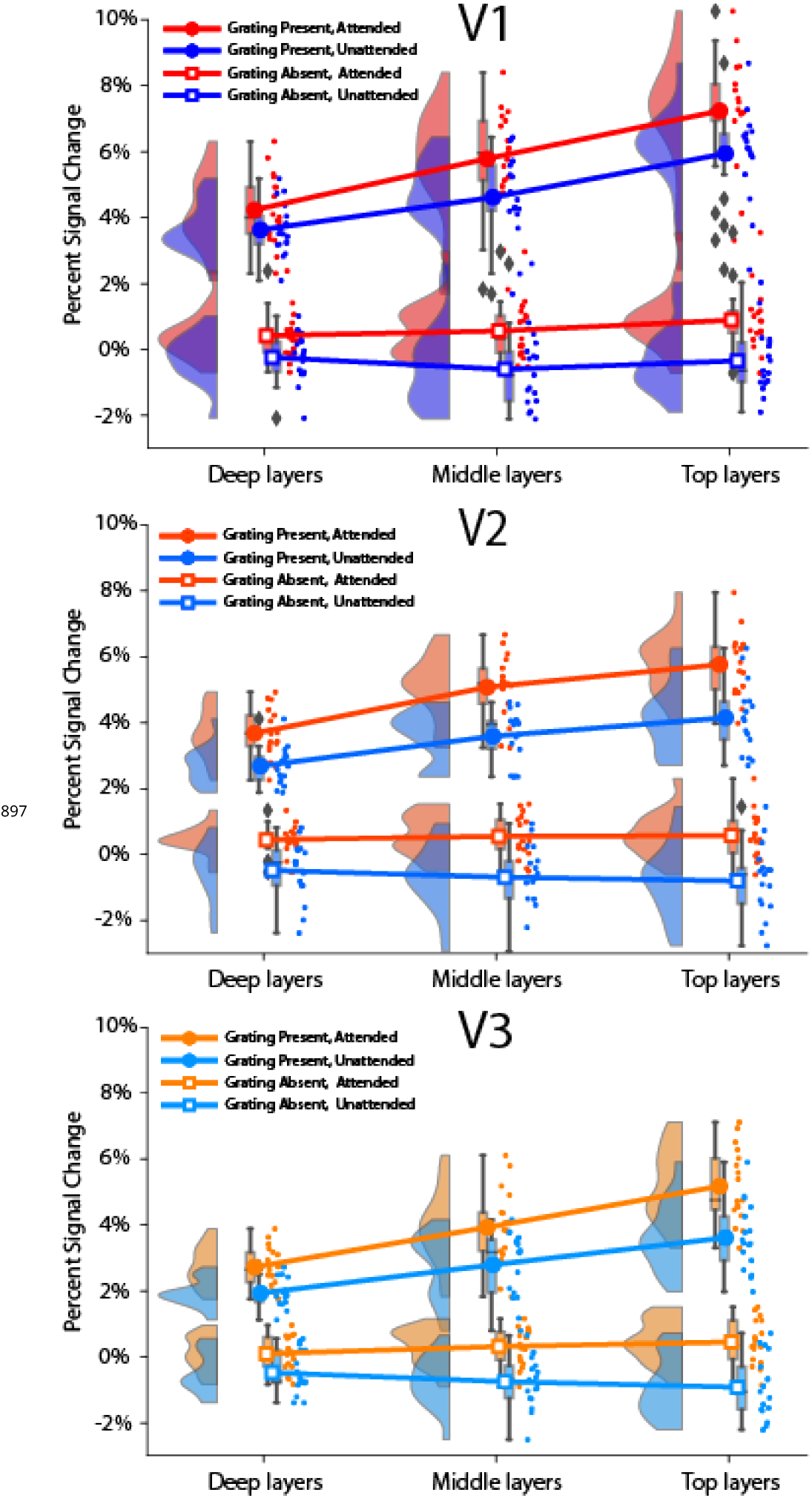
A control analysis that is identical to the main analysis, but repeated with a smaller region of interest. Only the 300 highest activated vertices were included in the analysis.

**Figure 3-Figure supplement 2.**
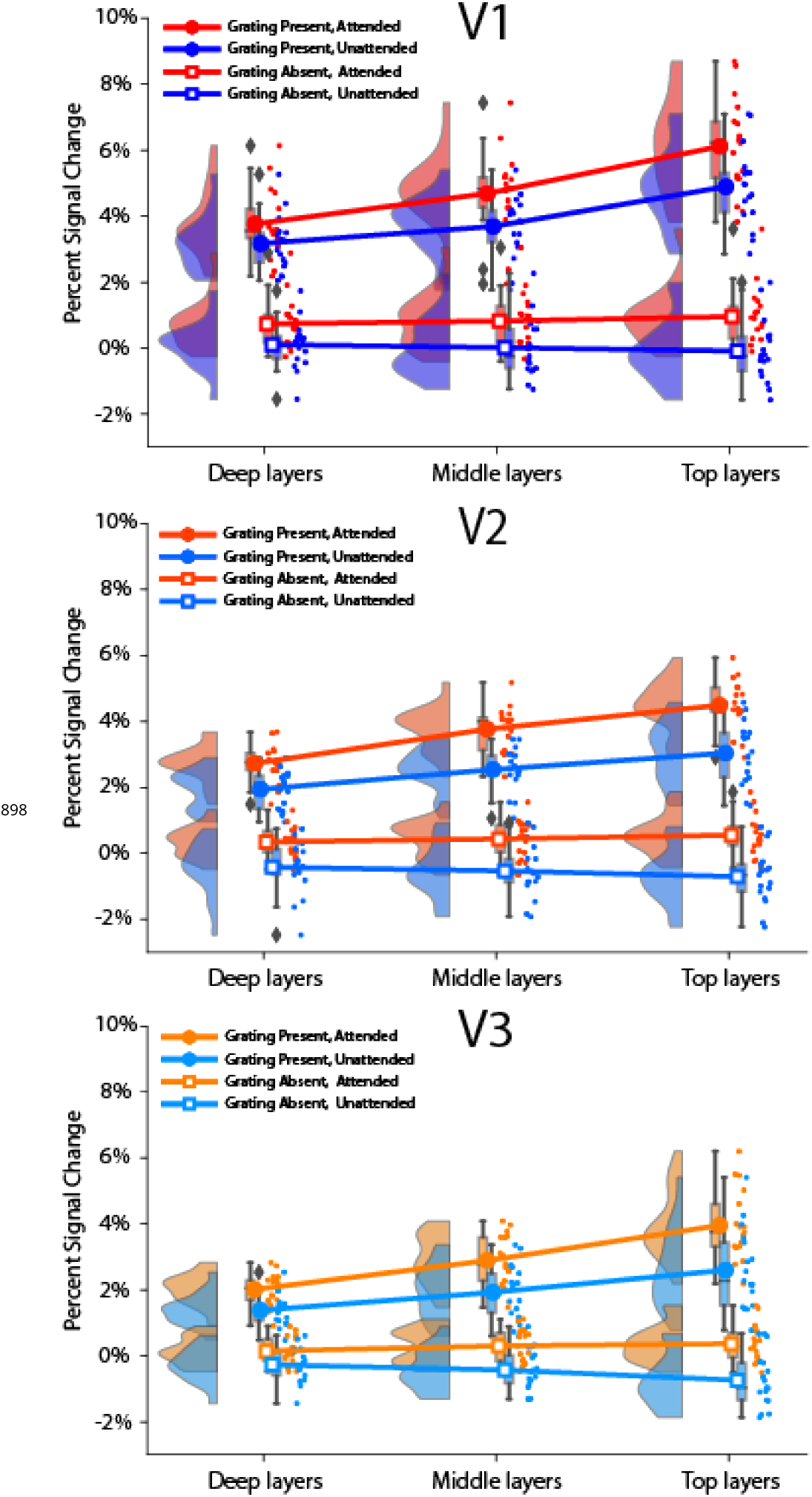
A control analysis that is identical to the main analysis, but repeated with a larger region of interest. Only the 900 highest activated vertices were included in the analysis.

**Figure 3-Figure supplement 3.**
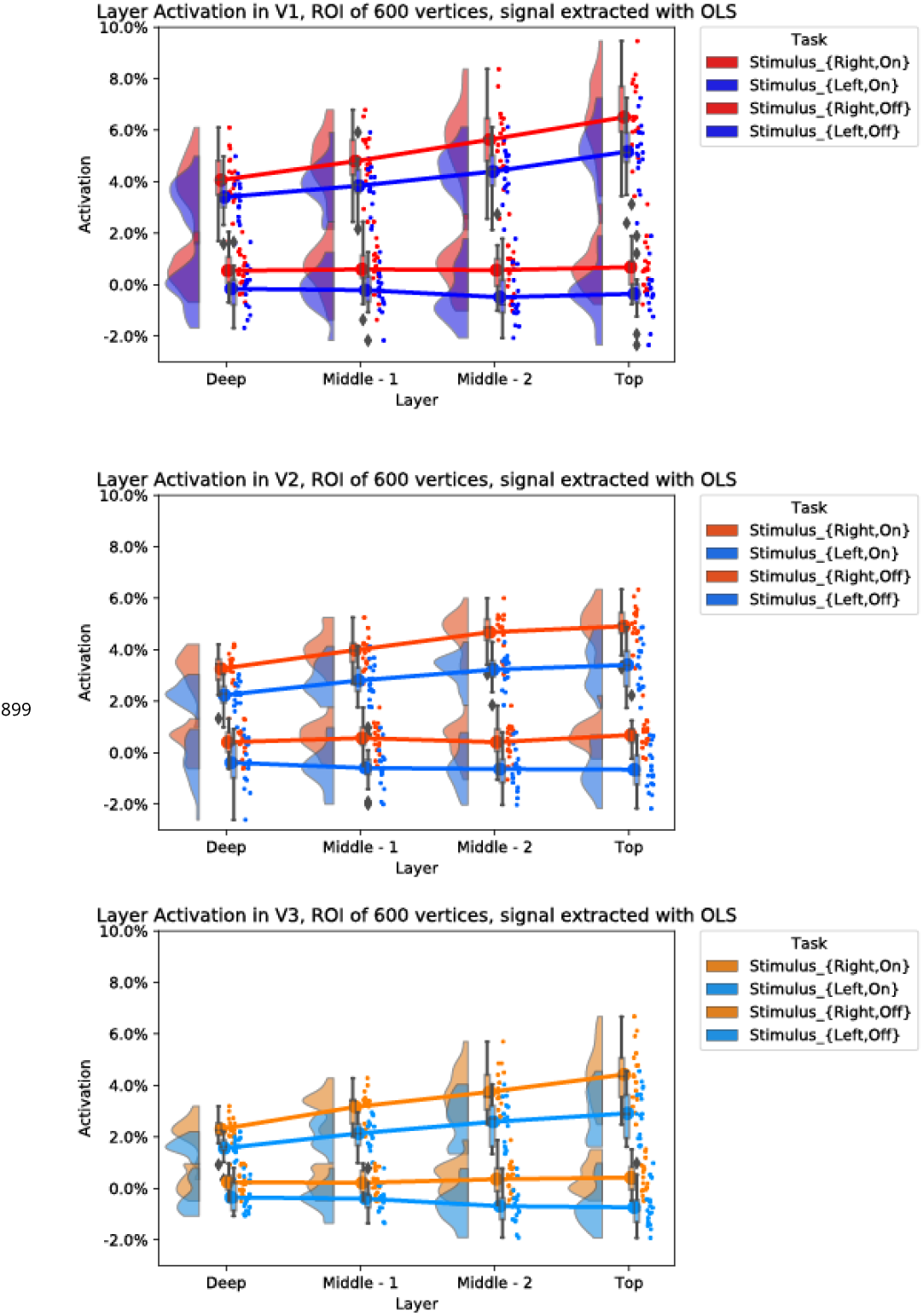
A control analysis that is identical to the main analysis, but repeated with four extracted layers instead of three.

**Figure 3-Figure supplement 4.**
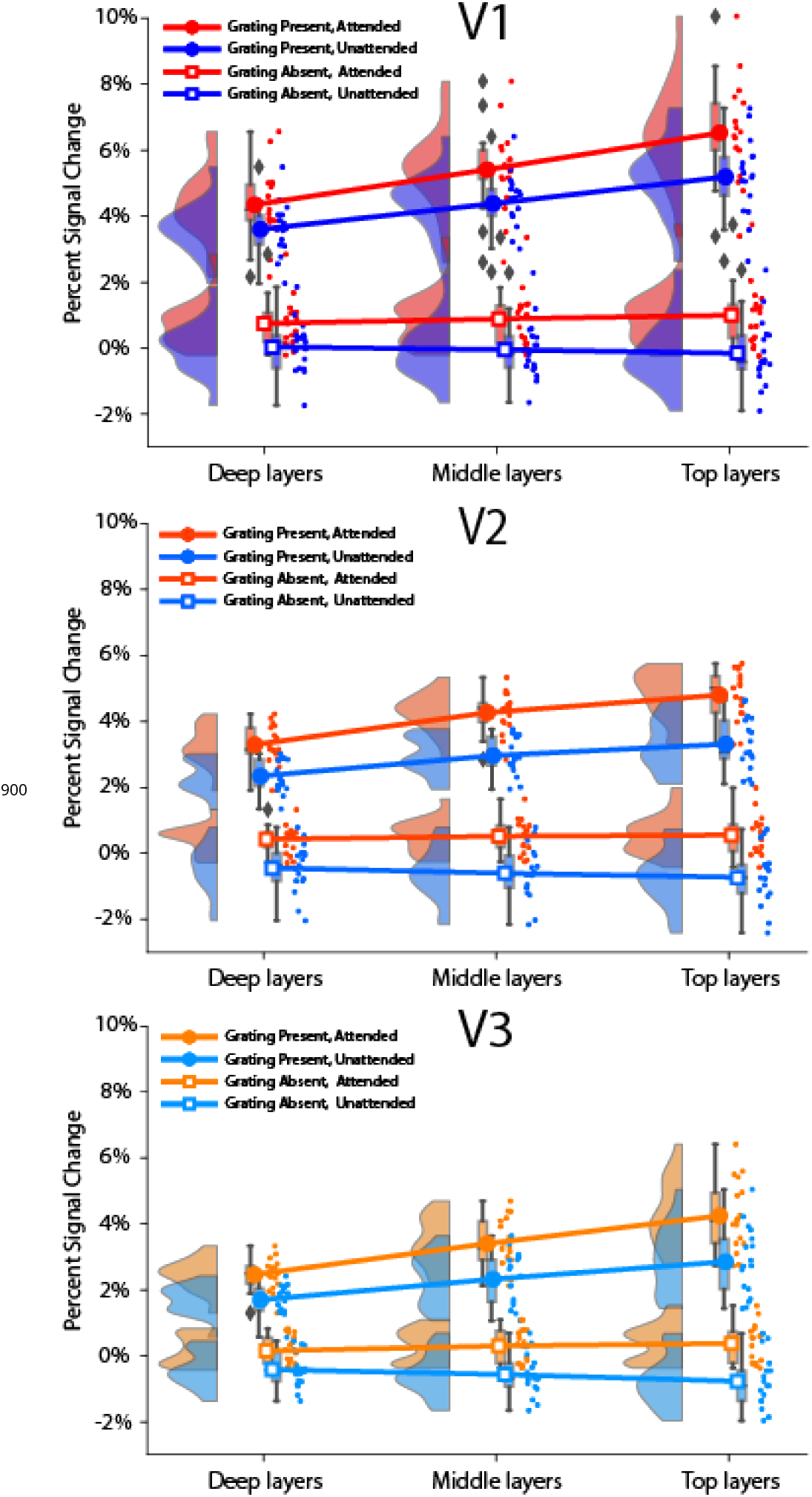
A control analysis that is identical to the main analysis, but repeated with a different way of obtaining laminar signal.

**Figure 4-Figure supplement 1.**
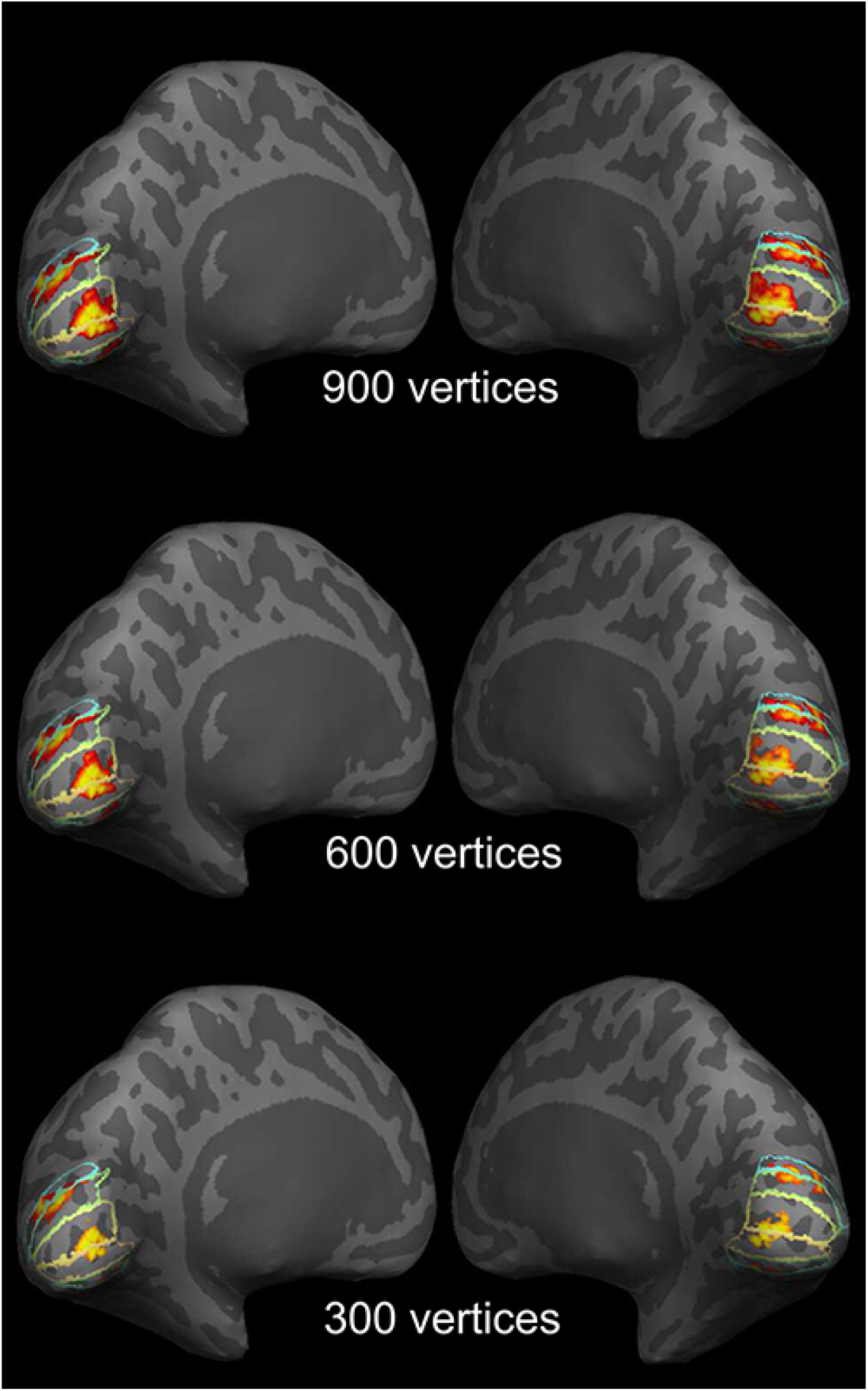
Example of Regions of Interest on the inflated cortical surface for a representative subject. The label contours from top to bottom show dorsal V3, V2, and V1 and ventral V1, V2, and V3, in both hemispheres. The 600 most activated vertices (coloured area) per region were selected for the main analysis. The 300 and 900 vertices were used for control analyses in order to show that the effects are independent of size of region of interest.

